# Contribution of B-1 cells in the cytotoxicity of CD8 T lymphocytes in encephalitozoonosis

**DOI:** 10.1101/2023.11.22.568300

**Authors:** Cristina Gabriela Nascimento de Oliveira, Elizabeth Cristina Perez, Anuska Marcelino Alvares-Saraiva, Maria Anete Lallo

## Abstract

*Encephalitozoon cuniculi* is an opportunistic intracellular pathogen that establishes a balanced relationship with immunocompetent individuals depending on the activity of their CD8^+^ T cells lymphocytes. However, lower resistance to experimental infection with *E. cuniculi* was found in B-1 deficient mice (Xid), besides increased the number of CD8 T lymphocytes. Here, we evaluated the cytotoxic activity of CD8^+^ T lymphocytes from Balb/c wild-type (WT) or Balb/c Xid mice (with B-1 cell deficiency) on the microbicidal activity of macrophages challenged with *E. cuniculi*. CD8 T lymphocytes from WT or Xid mice previously infected or not with *E cuniculi* were co-cultured with macrophages challenged with *E. cuniculi*. We evaluated macrophages viability and microbicidal activity, and proliferation, viability, and presence of activating molecules (CD62L, CD69, and CD107a) in CD8 T lymphocytes. Macrophages co-cultured with CD8 T lymphocytes from WT demonstrated high microbicidal activity. CD8 T lymphocytes obtained from uninfected WT mice had a higher proliferative capacity and a higher expression of CD69 and LAMP-1-activating molecules compared to Xid CD8^+^ T lymphocytes. CD8 T lymphocytes from infected Xid mice proliferated more than those from WT mice, however, when the expression of the activating molecule CD69 associated with the expression of CD62L was kept low. In conclusion, the absence of B-1 cells in Xid mice might be associated with the lower expression of activating molecules in CD8^+^ T lymphocytes and their cytotoxic activity. However, after a previous infection with *E. cuniculi*, CD8 T lymphocytes were more effective in killing macrophages infected with *E. cuniculi*.

## Introduction

Microsporidia are opportunistic, obligate intracellular parasites that affect vertebrate and invertebrate hosts [1]. The phylum Microsporidia has at least 200 genera and more than 1,500 species, of which 17 species affect mammals, including domestic animals and humans [2, 3] (Han et al., 2021; Mohamed et al., 2021). Microsporidian spores are resistant and infective; they are usually ingested by susceptible hosts and infect host cells by extruding their polar tubule, injecting the sporoplasm into the cytoplasm, which initiates their development and proliferation [4, 5]. To survive, the pathogen fights against the immune response using various adaptive evasion mechanisms. *Encephalitozoon cuniculi* is used as a standard for studying the immune response in mice [6]. It infects epithelial and endothelial cells, fibroblasts, and macrophages of various mammals, including rabbits, rodents, carnivores, monkeys, several wild mammals, and humans [6]. Several researchers have evaluated immunity against microsporidia, especially *E. cuniculi*, in recent years, and highlighted the importance of adaptive immunity in combating microsporidiosis [7, 8]. T cells protect against *E. cuniculi* infection [9, 10, 11]. Some studies have shown that protective immunity against intraperitoneal (i.p.) infection depends specifically on CD8 T cell immunity; TCD8^−/−^ mice died after inoculation of the pathogen, which caused severe and disseminated disease [12, 13, 14, 15]. In contrast, *E. cuniculi* infection of CD4^−/−^ mice did not result in any mortality [11]. Our group found that several immunosuppressive doses of cyclophosphamide can increase the susceptibility of mice to encephalitozoonosis by significantly reducing the population of T lymphocytes [16].

However, before acquired immunity is activated to control microsporidiosis, the innate immune response must play its role in antigen preparation and presentation and *milieu* modulation via the release of chemokines and cytokines, which subsequently activate adaptive immune cells. Macrophages, dendritic cells, natural killer cells (NKs), and innate-like lymphocytes are essential for eliciting responses against microsporidian infections [8]. Our group has investigated the effect of B-1 cells on encephalitozoonosis. B-1 cells are a part of innate immunity and constitute a distinct population from B (B-2) cells due to their anatomical location, phenotype, self-renewal capacity, and ability to produce natural antibodies [17]. Generally, B-1 cells are found in the pleural and peritoneal cavities and express high levels of IgM and low levels of IgD and CD11b on their surface [18]. B-1 cells play a key role in acquired and innate immunity. To assess their role, Balb/c Xid mice (Xid) are used because they are carriers of Bruton’s tyrosine kinase (Btk) deficiency, which leads to B cell deficiency [19, 20].

In another study, our group showed that Xid mice were more susceptible to infection by *E. cuniculi* than wild-type (WT) mice, via the oral and intraperitoneal route. This lower resistance was associated with B-1 cell deficiency since the adoptive transfer of these cells to Xid mice promoted the increase in immune cell populations and resulted in greater resistance to infection [21, 22]. We also showed that Xid mice treated with Cy immunosuppression developed severe and disseminated encephalitozoonosis, which was similar to that reported in the models with T-cell deficiency. However, populations of TCD8^+^ and TCD4^+^ lymphocytes in the peritoneum were higher than those in WT mice, which had milder disease [23]. B-1 cells influenced the composition of pulmonary granuloma induced by BCG in mice [24] and it was demonstrated that B-1 cells induced a large infiltrate of CD8+ T lymphocytes in grafts [25]. Additionally, it has been suggested that peritoneal B-1 cells may also influence the immune response by activating T cells without necessarily migrating to the lymphoid organ responsible for eliciting a response [26].

We hypothesized that partial susceptibility of Xid mice to an infection may suggest an inefficient function of T lymphocytes due to the absence of B-1 cells. Our aim was to study the effects of CD8^+^ T lymphocytes obtained from WT mice (with B-1 cells) and Xid mice (with B-1 cell deficiency) on the microbicidal activity of macrophages challenged with *E. cuniculi*.

## Methods

### Ethics statement

All experiments involving animals were performed following the recommendations outlined in the Guide for the Care and Use of Laboratory Animals (Concea – Conselho Nacional de Controlee m Experimentação Animal). All research protocols for animals (number 016/17) were approved by the institutional animal use committee (Comissão de Ética no Uso de Animais – CEUA) at Universidade Paulista - UNIP. We prioritized minimizing the suffering and distress of animals during the study.

### Encephalitozoon cuniculi spores

The spores of *E. cuniculi* (genotype I) (Waterborne Inc., New Orleans, LA, USA) used in this study were previously cultivated in a rabbit kidney cell lineage (RK-13, ATCC CCL-37) in RPMI medium supplemented with 10% fetal calf serum (FCS-Sigma-Aldrich, St. Louis, MO, USA) and gentamicin at 37 °C in a humidified atmosphere with 5% CO_2_. The spores were purified by centrifugation at 500 *g* for 20 min, and cellular debris was excluded using 50% Percoll. The spores of *E. cuniculi* were counted using a Neubauer chamber.

### Animals

Isogenic, female, eight-week-old, and specific pathogen-free Balb/c WT and Xid mice were obtained from the “Centro de Desenvolvimento de Modelos Experimentais para Biologia e Medicina” (CEDEME) at Federal University of Sao Paulo (UNIFESP, in Portuguese). The animals were placed in groups (uninfected or infected) and kept in sterilized individual cages at the animal facility at Paulista University (UNIP), under controlled conditions of temperature and humidity. All animals were provided water and food *ad libitum*.

### Macrophage culture and infection

We maintained RAW 264.7 macrophages in RPMI medium (Sigma-Aldrich, St Louis, MO, USA) supplemented with 10% FCS (R10) at 37 °C in a humidified atmosphere with 5% CO_2_. The cultures were maintained by periodically changing the R10 medium to obtain macrophages. The medium was changed every two days until 70 to 80% confluence was reached.

Macrophages were inoculated in 24-well plates (1×10^5^ cells/well) in 300 µL of R10 medium and incubated for 4 h at 37 °C in an atmosphere containing 5% CO_2_. After removing the supernatant, the cultures were infected with *E. cuniculi* spores at a 2:1 ratio of *E. cuniculi* spores to each type of macrophage for 1 h. The experiments were performed in quadruplicates. Before being co-cultured with CD8 T lymphocytes, the supernatant of macrophages was removed, and the wells were washed with PBS to remove spores and cellular debris.

### Isolation of CD8 T lymphocytes

The CD8^+^ T cells from the splenocyte population were sorted by flow cytometry FACS ARIA II at the Immunology Discipline of the Department of Microbiology and Immunology of the Federal University of São Paulo (UNIFESP). WT or Xid mice infected or not infected were administered ketamine (100 mg/mL), xylazine (20 mg/mL), and fentanyl (0.05 mg/mL) intraperitoneally. After cardiorespiratory arrest, the animals were washed with a chlorhexine solution and immersed in 70% alcohol. The spleens were collected under sterile conditions and mechanically dissociated using a syringe plunger and a 70 µm nylon mesh (cell strainer) in a PBS solution supplemented with 2% FBS (PBS+FBS). After centrifugation, the supernatant was discarded, and 1 mL of hemolytic buffer was added. After 2 min, 10 mL of PBS+FBS was added, and the samples were centrifuged at 1,500 rpm for 5 min. Next, the samples were incubated for 20 min with anti-CD16/CD32 antibodies to block Fc receptors under refrigeration. The cells were washed with PBS+FBS and incubated with the monoclonal antibody anti-mouse CD8 T conjugated to Fluorescein Isothiocyanate (FITC) (BD-Pharmingen, San Diego, CA), which was suitable for determining the phenotype of CD8^+^ T cells. After the cells were refrigerated for 20 min, they were washed and resuspended in 2 mL of PBS for cell sorting using a flow cytometer (Supplementary figure).

### Experimental Design

To understand the effects of B-1 cells on CD8 T lymphocytes, two experimental protocols were used. Using the first protocol, naïve CD8^+^ T lymphocytes were collected from the spleen of uninfected WT and Xid mice and separated by cell sorting. Using the second protocol, primed CD8^+^ T lymphocytes were collected by cell sorting from WT and Xid mice after 14 days of infection with 1×10^7^ spores of *E. cuniculi*, by i.p. route. In both protocols, CD8 T cells were co-cultured with *E. cuniculi*-infected macrophages for 36 h. After incubation, we analyzed macrophages viability and their microbicidal potential against *E. cuniculi* by measuring the proliferative capacity of the recovered spores to develop in RK cell culture. For CD8 T lymphocytes, we evaluated the viability, proliferative capacity, and activation profile of following the two protocols (Fig. 1).

**Figure 1.**
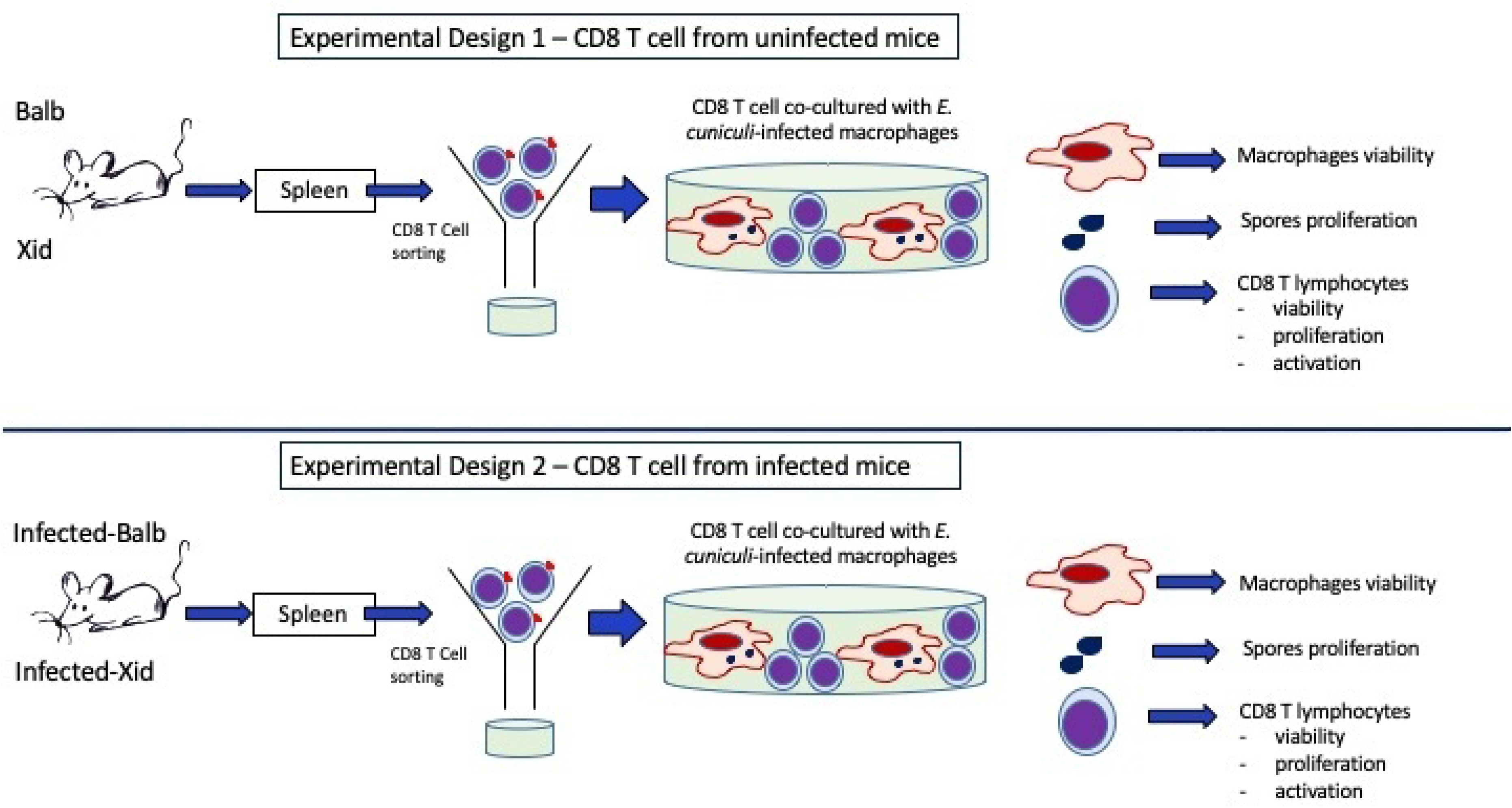
Graphical representation of the experimental strategies. <u>Experimental design 1</u> - CD8^+^ T lymphocytes were collected from the spleens of Balb/c or WT mice (with B-1 cells) and Xid (with B-1 cell deficiency). They were separated by cell sorting and co-cultured with macrophages previously infected with spores of *Encephalitozoon cuniculi*. After 36 h, the macrophages were tested for viability, the spores were tested for proliferation, and CD8 T cells were analyzed for viability, proliferation, and panel activation. <u>Experimental design 2</u> - CD8^+^ T lymphocytes were collected from the spleens of Balb/c or WT mice (with B-1 cells) and Xid (with B-1 cell deficiency) infected with *E. cuniculi* spores. They were separated by cell sorting and co-cultured with macrophages previously infected with the spores of *E. cuniculi*. After 36 h, the macrophages were tested for viability, the spores were tested for proliferation, and CD8 T cells were analyzed for viability, proliferation, and panel activation.

### Evaluation of the phagocytosis of spores by fluorescence microscopy

Macrophages were labeled with the fluorescent dye PKH-26 (Sigma Aldrich, St. Louis, MO, USA) according to the manufacturer’s instructions. Briefly, after diluent C and fluorophore were added to cells, the mixture was incubated for 5 min. After blocking with 5% FBS, the cells were washed and cultured in 24-well plates on glass coverslips for 4 h. Then, macrophages were infected with spores of *E. cuniculi* labeled with the fluorescent dye carboxyfluorescein succinimidyl ester (CFSE) in a 2:1 ratio and incubated for 1 h. In different groups, lymphocytes from mice (WT or Xid, both uninfected) were added to macrophages culture, according to the experimental design, and incubated for 36 h at 37°C with 5% CO_2_. As a negative control, *E. cuniculi* infected macrophages were labeled. After incubation, the coverslips were washed with PBS and fixed with 300 µL of 4% paraformaldehyde for 20 min. Then, the coverslips were washed with PBS and mounted on a glass slide using 5 µL of the mounting medium DAPI-Fluorishield (Sigma Aldrich, St. Louis, MO, USA). All observations were performed using a Leica TCS SP8 confocal microscope, and the LAS X program was used for analysis.

### Viability of macrophages and CD8 T cells

Viability was analyzed with Kit 7AAD/Annexin (BD Pharmingen), according to the manufacturer’s instructions. Before viability assay, the nonspecific Fc receptors from macrophages or CD8 T cells were blocked with anti-CD16/32 antibodies and incubated for 15 min. After washing with Mac’s buffer (0.5% BSA and 0.075% EDTA), the macrophages were labeled with mouse anti-F4/80 monoclonal antibody conjugated to phycoerythrin (PE) and CD8 T cells with FITC-conjugated mouse anti-CD8 monoclonal antibody for 30 min at 4 °C. Once again, the washing with Mac’s buffer was performed. After that, the cells were centrifuged and resuspended in the Kit buffer, followed by the addition of 1 µL of annexin and 1 µL of 7AAD. After 15 min of incubation at room temperature in the dark, the kit buffer was again added, and apoptosis quantification was performed on an Accuri^TM^C6 flow cytometer (BD Biosciences, Mountain View, CA) while collecting after 30 s from each of the tubes.

### Lymphocyte proliferation assay

The CD8^+^ T lymphocytes from the spleens of WT and Xid mice (infected and uninfected) were isolated and labeled with CFSE, following the manufacturer’s instructions. Briefly, 1 µM CFSE was added to the lymphocytes and incubated at 37 °C for 10 min. Then, two washes were performed with sterilized PBS. CFSE-CD8 T lymphocytes were co-cultured with macrophages infected with *E. cuniculi* spores (described in Section 2.5) for 36 h at 37 °C with 5% CO_2_. After incubation, the supernatant containing the lymphocytes was collected, centrifuged, and resuspended in 150 µL of PBS for analyzing the samples using a flow cytometer (ACCURI C6 Sampler; BD Bioscience); the decrease in the CFSE fluorescence intensity was used as a proliferation parameter.

### Analysis of the cytotoxic activity

Macrophages infected with *E. cuniculi* (details in Section 2.5) were co-cultured with CD8^+^ T lymphocytes from WT or Xid mice, previously separated by sorting, in a 5:1 ratio (five lymphocytes to one macrophage). The cells were incubated at 37 °C with 5% CO_2_. After 31 h, 10 µL of mouse anti-CD107a (LAMP1) monoclonal antibodies conjugated to Phycoerythrin (PE) were added per well. After 1 h, 0.5 µL of monensin and 0.5 µL of brefeldin were added to the final solution (500 µL) to block protein transport across the cell membrane. Upon completion of the experiment after 36 h, the supernatant was collected, the adhered macrophages were released using a cell scraper, and they were added to the lymphocytes in the supernatant. After centrifugation at 200g for 5 minutes, the anti-Fc receptors were added to cells, with the subsequent separation of the sediment into two aliquots. In panel 1, the markers used included FITC-conjugated mouse anti-CD8 monoclonal antibody, allophycocyanin (APC)-conjugated mouse anti-CD69 monoclonal antibody, and Peridinin Chlorophyll Protein Complex (PerCP)-conjugated mouse monoclonal antibody (Cy5). The cells were incubated for 30 min under refrigeration and away from light, washed and resuspended in 150 µL of PBS, and analyzed using a flow cytometer. In panel 2, lymphocytes were labeled using interferon-gamma (IFN-γ). Briefly, the samples were permeabilized with a Fixation/Permeabilization solution (BD Citofix/Citoperm - Fixation/Permeabilization kit) for 20 min and then washed with Perm/Wash buffer. Next, the cells were incubated for 30 min with FITC-conjugated anti-mouse CD8, PerCP-conjugated anti-F4/80, and APC-conjugated anti-IFN-γ antibodies. Finally, the cells were washed with Perm/Wash buffer, resuspended in 150 µL of PBS, and analyzed using an Accuri BD flow cytometer.

### Microbicidal activity

Macrophages infected with *E. cuniculi* spores at a 2:1 ratio were co-cultured with CD8^+^ T lymphocytes from WT or Xid mice, at a lymphocyte-to-macrophage ratio of 5:1 under controlled conditions (37 °C and 5% CO_2_). After 36 h, both types of cells were harvested and disrupted by alternately exposing them to cold (ice bath) and heat (70 °C) treatment for 20 s in each condition. The process was repeated 20 times for the complete disruption of cells and release of intracellular spores. The total number of spores collected was denoted as T0. To determine that the spores were viable, they were inoculated into RK-13 cells and cultured for seven days. Following this, the cells were harvested and disrupted, and the number of spores present (T7) was counted. The T7/T0 ratio (Ratio = final number of T7 spores/initial number of T0 spores) was evaluated and indicated the proliferation of spores. The averages of the ratios of each group were used to conduct the statistical analysis.

### Statistical analysis

All differences between groups were determined by paired t-tests. All values were presented as the mean ±standard error of the mean (SEM) with a significance of α = 0.05 (p < 0.05). All graphs were plotted using “GraphPad Prism” version 5.0 for Windows^®^ (GraphPad Software Inc., La Jolla, CA, EUA).

## Results

### Co-cultures of infected macrophages and CD8 T lymphocytes

The spores, with oval shape and fluorescence green color, were internalized by macrophages after 1 h of observation (Fig. 2A). However, after 36 h of incubation, in co-cultures with CD8 T lymphocytes from WT and Xid mice, few spores were detected with their characteristics preserved (Fig. 2B and 2C). Also, fewer intact spores with fluorescence green were detected inside the macrophages. However, the presence of orange vacuoles indicated that phagolysosomes were frequent, suggesting spore degradation (Fig. 2B and 2C). The contact between CD8 T lymphocytes and macrophages was also visible, indicating intercellular communication occurred (Fig. 2C).

**Figure 2.**
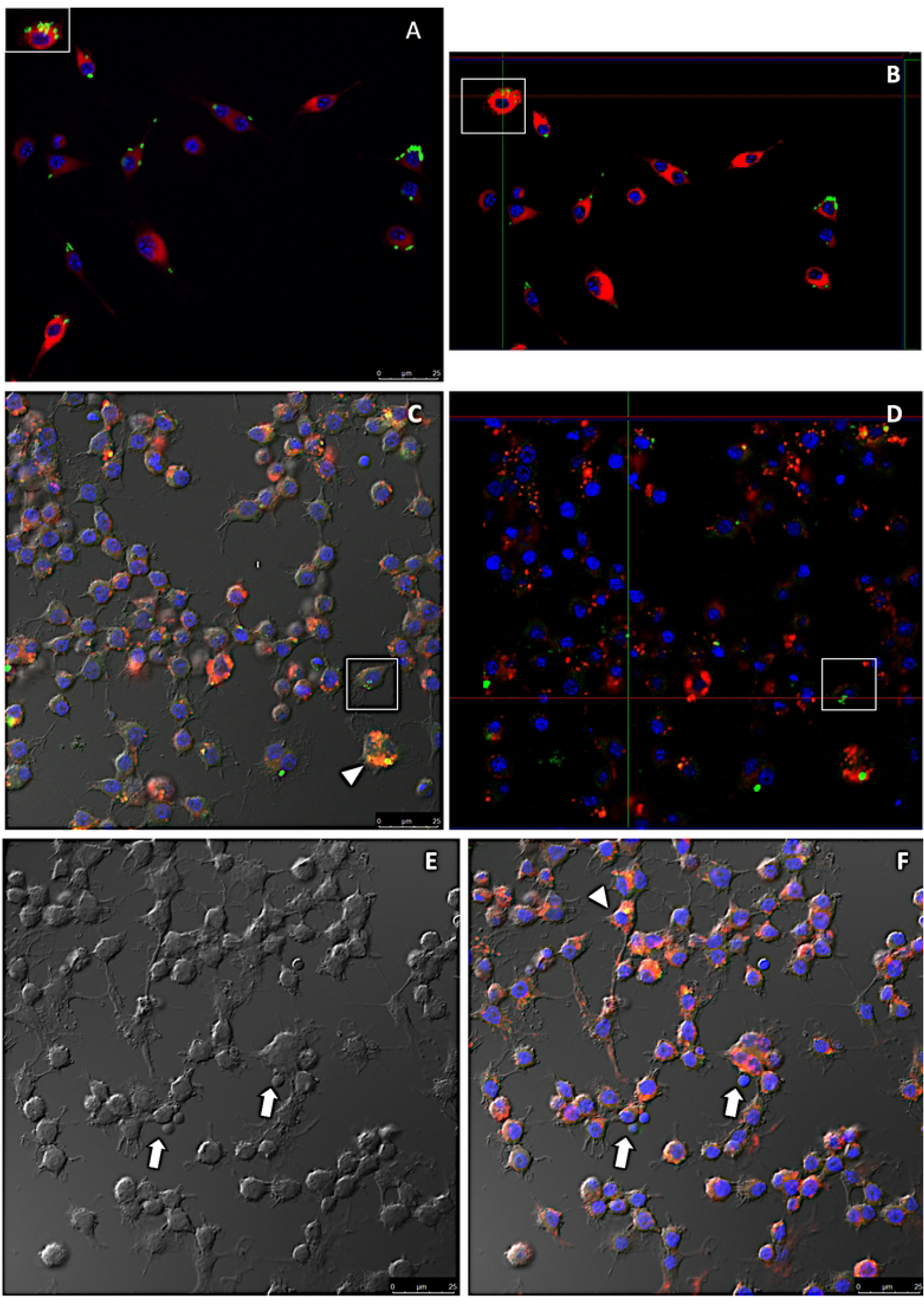
Photomicrographs of *E. cuniculi* infected macrophages were or were not co-cultured with CD8 T lymphocytes, obtained by confocal microscopy. Macrophages incorporated with the PKH-26 dye (red) were infected with *E. cuniculi* spores, stained with CFSE (green) for 1 h, and the slides were mounted with DAPI (blue). Macrophages incorporated with the PKH-26 dye (red) were infected with CFSE-stained *E. cuniculi* spores (green) for 1 h, and the slides were mounted with DAPI (blue) Note the presence of *E. cuniculi* spores inside macrophages after 1 h of infection (insert). (A) Macrophages incorporated with the PKH-26 dye (red) were infected with the spores of *E. cuniculi* stained with CFSE (green) for 1 h, and the slides were mounted with DAPI (blue). Then, CD8^+^ T lymphocytes from WT mice were co-cultured with macrophages for 36 h. (C) Macrophages incorporated with the PKH-26 dye (red) were infected with the spores of *E. cuniculi* stained with CFSE (green) for 1 h, and the slides were mounted with DAPI (blue). Then, CD8^+^ T lymphocytes from Xid mice were co-cultured with macrophages for 36 h. Few intact spores were present inside macrophages. CD8 T lymphocytes can be seen adhering to macrophages (arrow).

### Previous infection with *E. cuniculi* reduced the viability of macrophages

The viability of macrophages co-cultured with CD8 T lymphocytes obtained from uninfected WT and Xid mice was similar; around 90–95% of the cells were viable (Fig. 3A, 3B). Death due to apoptosis was twice as high as that due to necrosis in the two groups of cells. However, the viability of macrophages co-cultured with CD8 T lymphocytes from *E. cuniculi* infected Xid mice previously was lower (60–70%) (Fig. 3E, 3F) and most cells died due to apoptosis (Fig. 3G, 3H). These results suggested that lymphocytes from Xid mice infected with *E. cuniculi* induced an increase in the apoptosis of macrophages.

**Figure 3.**
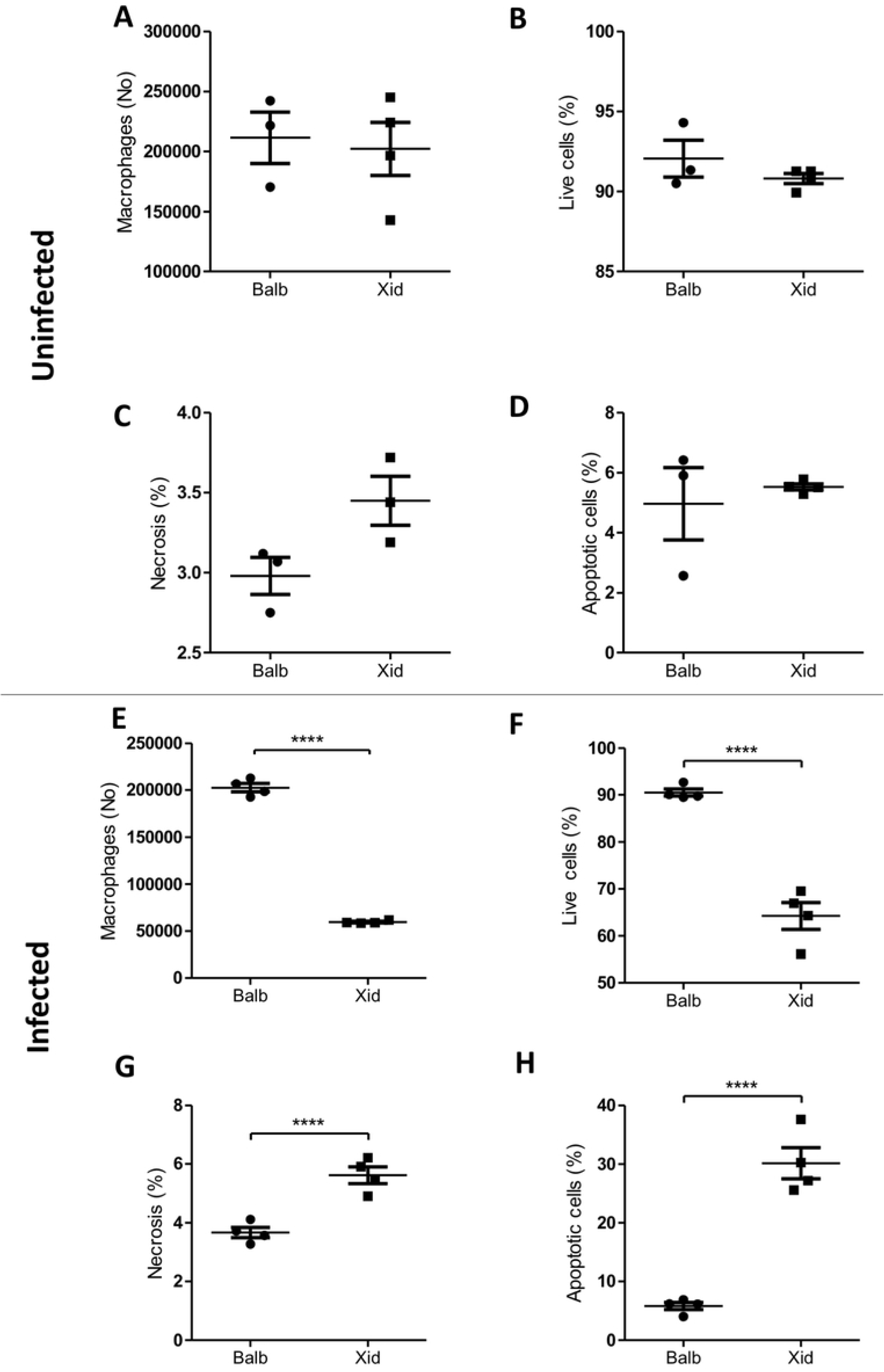
Viability analysis of CD8 T lymphocytes co-cultured with *E. cuniculi*-challenged macrophages. The viability of CD8 T lymphocytes from uninfected mice or those previously infected with *E. cuniculi* was determined after co-culture with *E. cuniculi*-challenged macrophages. (A-D) Macrophages challenged with *E. cuniculi* were co-cultured for 36 h with CD8 T lymphocytes collected from uninfected Balb/c (WT) and Xid mice. (E-H) Macrophages challenged with *E. cuniculi* were co-cultured for 36 h with CD8 T lymphocytes collected from *E. cuniculi*-infected Balb/c (WT) and Xid mice. (A) The number of CD8 T lymphocytes. (B) The percentage of living CD8 T lymphocytes. (C) The percentage of CD8 T lymphocytes undergoing necrosis. (D) The percentage of CD8 T lymphocytes undergoing apoptosis. (E) The number of CD8 T lymphocytes. (F) The percentage of living CD8 T lymphocytes. (G) The percentage of CD8 T lymphocytes undergoing necrosis. (H) The percentage of CD8 T lymphocytes undergoing apoptosis. Statistical analysis: paired t-tests were conducted in triplicate; *** p < 0.001 and **** p < 0.0001.

The spores recovered from the co-cultures of macrophages and CD8 T cells were inoculated into RK cells. As *E. cuniculi* spores have a high affinity for RK cells, viable spores could internalize the sporoplasm through the polar tubule and multiply inside these cells. The results of the proliferation assay showed that the spores recovered from co-cultures of macrophages with CD8 T lymphocytes from uninfected WT mice were not viable (Fig. 4A), as the ratio between the spores inoculated in the RK cells to those recovered was zero. This finding showed the high microbicidal capacity of the macrophages in co-culture with CD8 T cells from WT uninfected mice. The proliferation rate of spores recovered from co-cultures of macrophages with CD8 T lymphocytes from uninfected Xid mice was high, i.e., the number of spores recovered from RK cell cultures was 2–10 times greater than the number of spores initially inoculated, these results confirmed the viability and subsequent multiplication of the spores in RK cells (Fig. 4A). Our findings indicated a decrease in the microbicidal activity of macrophages that were co-cultured with CD8 T lymphocytes from Xid mice.

**Figure 4.**
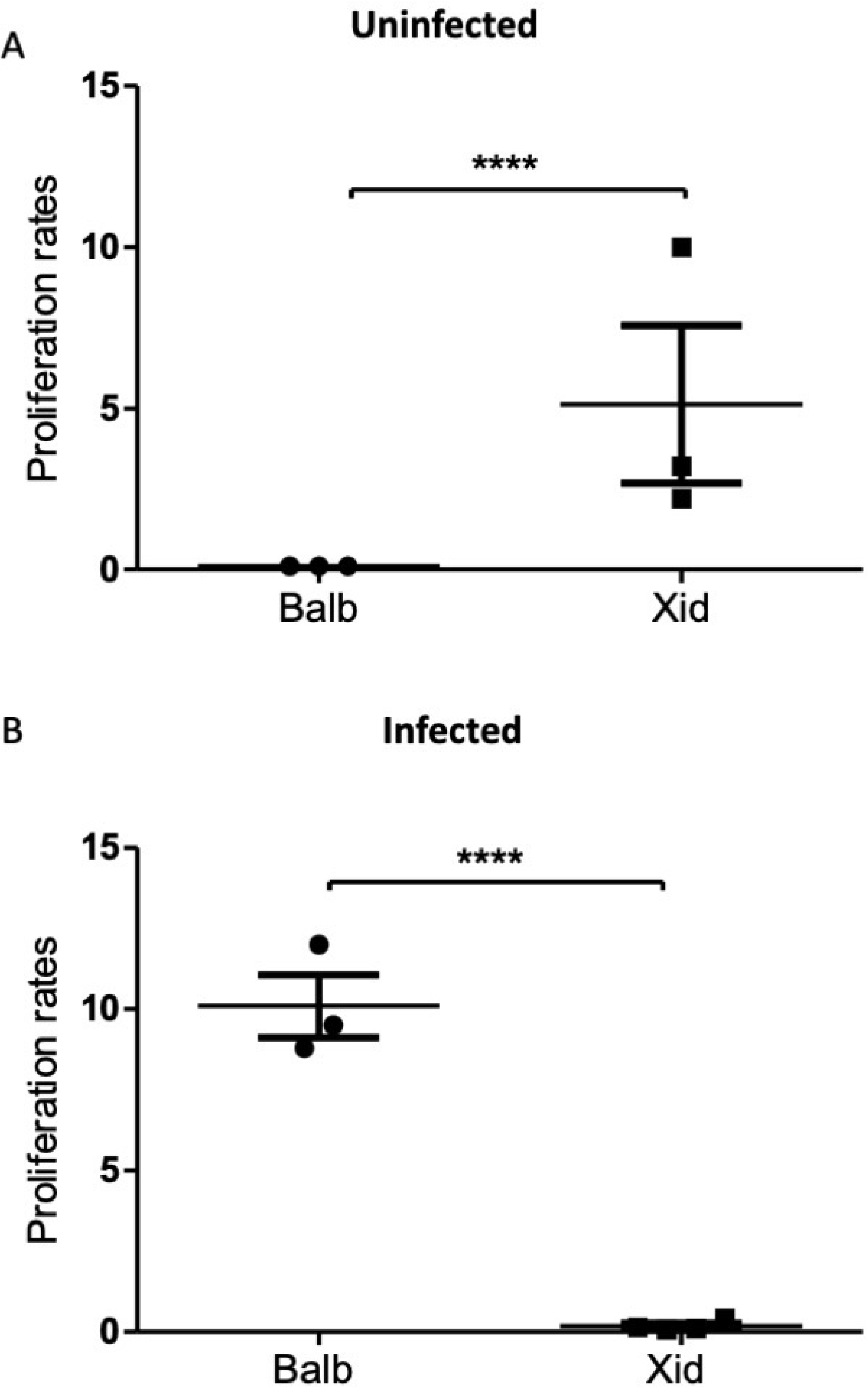
Proliferation ratio of spores obtained from macrophages co-cultured with CD8 T lymphocytes. After macrophages were co-cultured with CD8 T lymphocytes from Balb (WT) or Xid (uninfected or infected) mice, the spores recovered from cultures with macrophages challenged with *E. cuniculi* were counted (T0) and cultured in RK cells for seven days (T7) to obtain the ratio of proliferation (T7/T0). (A) CD8 T lymphocytes from uninfected Balb or Xid mice (Uninfected) were co-cultured with macrophages and challenged with *E. cuniculi*. (B) CD8 T lymphocytes from Balb or Xid mice previously infected with *E. cuniculi* (Infected) were co-cultured with macrophages and challenged with *E. cuniculi*. Statistical analysis: paired t-tests were conducted in triplicate; *** p < 0.001 and **** p < 0.0001.

The results of the proliferation assay performed for spores obtained from co-cultures with lymphocytes from infected WT and Xid mice showed the opposite pattern. A higher microbicidal activity was recorded in co-cultures with CD8 T lymphocytes from Xid mice (Fig. 4B).

### CD8 T lymphocytes from Infected Xid mice showed low viability and proliferation

The viability of CD8 T lymphocytes from WT or Xid uninfected was greater than 90%. (Fig. 5) The highest percentage of live CD8 T cells was recorded in cultured Xid mice (Fig. 5B). We observed a predominance of necrosis with a type of cell death associated with a decrease in CD8 T lymphocytes, WT vs Xid (Fig. 5C). CD8 T lymphocytes obtained from previously infected WT or Xid in coculture with macrophages demonstrated lower viability, between 75 and 85%, with the percentage of necrosis being 14% to 16% and 18 to 22%, observed respectively in WT and Xid (Fig. 5E, 5F, 5G), although the percentage of apoptosis was higher (2%) in lymphocytes from the mice in the Xid group (Fig. 5H).

**Figure 5.**
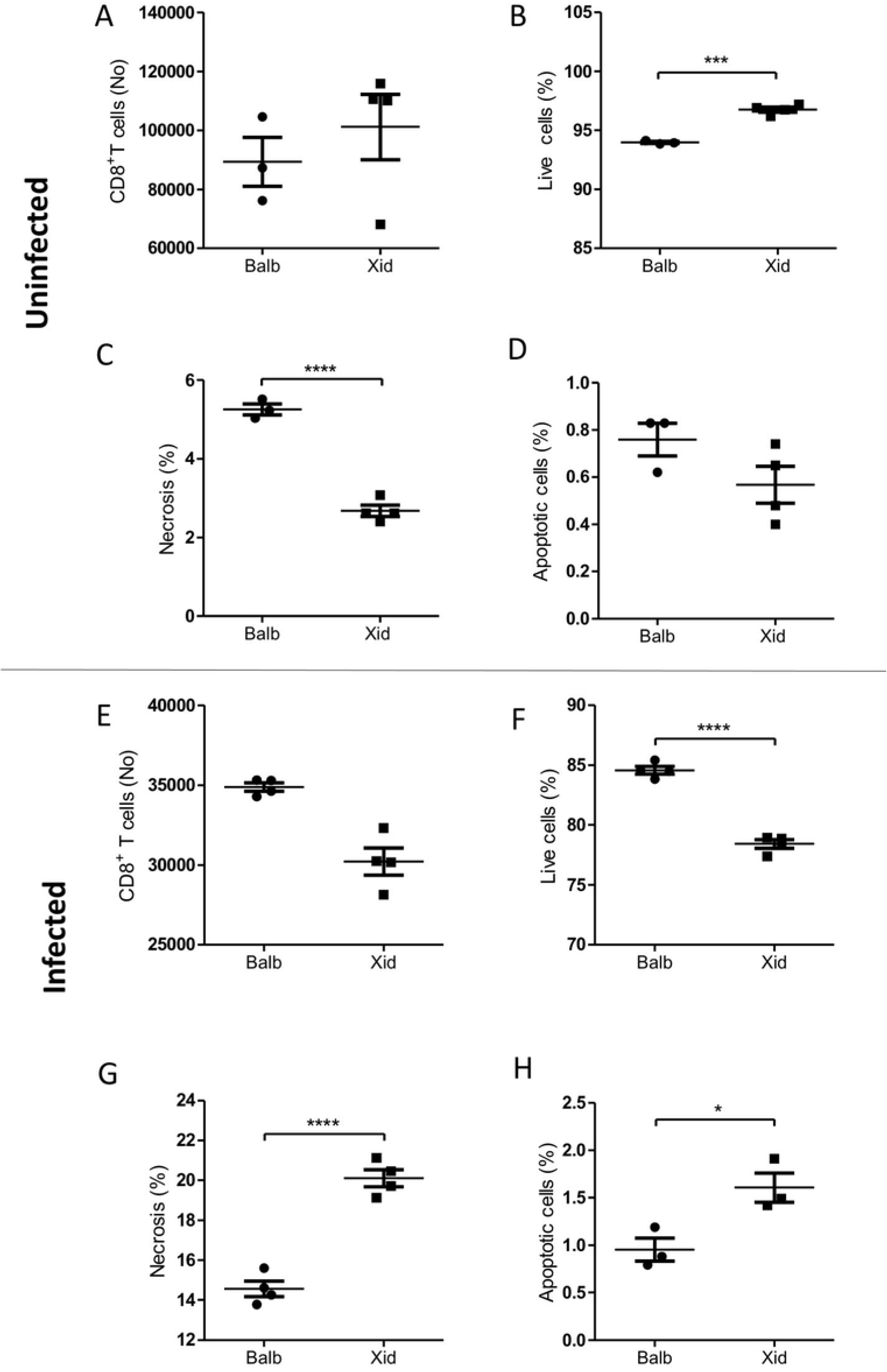
Viability analysis of *E. cuniculi*-challenged macrophages co-cultured with primed or unprimed CD8 T lymphocytes. (A-D) Macrophages challenged with *E. cuniculi* were co-cultured for 36 h with CD8 T lymphocytes collected from uninfected Balb/c (WT) mice and Xid mice. (E-H) Macrophages challenged with *E. cuniculi* were co-cultured for 36 h with CD8 T lymphocytes collected from *E. cuniculi*-infected Balb/c (WT) mice and Xid mice. (A) The number of macrophages. (B) The percentage of living macrophages. (C) The percentage of macrophages undergoing necrosis. (D) The percentage of macrophages undergoing apoptosis. (E) The number of macrophages. (F) The percentage of living macrophages. (G) The percentage of macrophages undergoing necrosis. (H) The percentage of macrophages undergoing apoptosis. Statistical analysis: paired t-tests were conducted in triplicate; * p < 0.05 and **** p < 0.0001.

The proliferation rate was higher in CD8^+^ T lymphocytes from uninfected WT mice compared to that of the CD8^+^ T lymphocytes from other experimental groups. This finding was confirmed by the low mean CFSE fluorescence intensity (MIF CFSE) observed in this group (Fig. 6). Our results suggested that CD8 T lymphocytes from Xid mice showed low proliferation irrespective of the presence of microsporidia. In infected WT mice, the number of CFSE low lymphocytes from infected mice was much lower and the presence of the pathogen probably decreased cell proliferation as well as increased death from necrosis.

**Figure 6.**
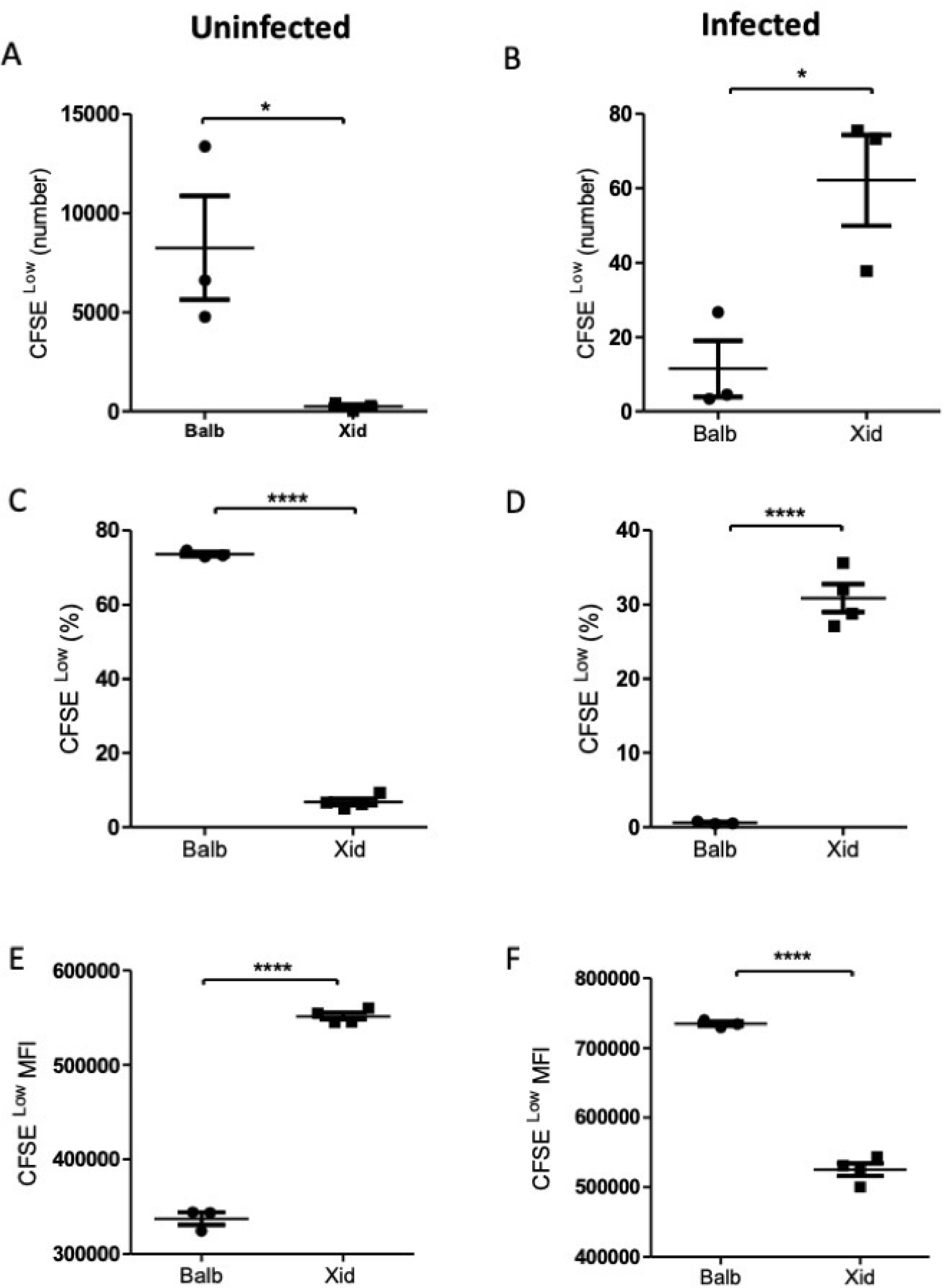
Evaluation of the proliferation of CD8 T lymphocytes. Analysis of the percentage, number, and mean fluorescence (MFI) of CFSE incorporated into lymphocytes immediately after collection by separation and sorting. (A) The number of CD8 T lymphocytes CFSE^l^°^w^ from uninfected Balb/c or Xid mice. (B) The number of CD8 T lymphocytes CFSE^l^°^w^ from Balb/c or Xid mice infected with *E. cuniculi*. (C) The percentage of CD8 T lymphocytes CFSE^l^°^w^ from uninfected Balb/c or Xid mice. (D) The percentage of CD8 T lymphocytes CFSE^l^°^w^ from infected Balb/c or Xid mice. E) The mean fluorescence intensity (MIF) of CD8 T lymphocytes CFSE^l^°^w^ from uninfected Balb/c or Xid mice. (F) The mean fluorescence intensity (MIF) of CD8 T lymphocytes CFSE^l^°^w^ from infected Balb/c or Xid mice. Statistical analysis: paired t-tests were conducted in triplicate; *p < 0.05 and **** p < 0.0001.

### CD8 T cells from uninfected WT mice with high expression of CD69 and Lamp-1 activation increased the microbicidal action of macrophages

The expression of CD69 selectin was higher in CD8 T lymphocytes from WT mice compared to that in the lymphocytes from Xid mice, in the uninfected and infected groups (Fig. 7A, 7B). These results suggested that B-1 cells, abundant in WT mice and deficient in Xid mice, probably influenced the expression of co-stimulatory molecules, such as CD69 in CD8 T lymphocytes. These findings explained the lower resistance of Xid mice to encephalitozoonosis described by our group (Langanke et al., 2017). We found that the expression of Lamp-1 increased in the lymphocytes of uninfected WT mice (Fig. 7A, 7B). We examined the correlation between the co-stimulatory molecules CD62 x CD107 (Lamp-1) and CD62 x CD69 and found an inversely proportional relationship, i.e., the expression of CD62L decreased as the expression of CD107 or CD69 increased on the surface of CD8 T lymphocytes. For the first time, we illustrated the dynamics of the expression of activation molecules on the surface of lymphocytes in encephalitozoonosis.

**Figure 7.**
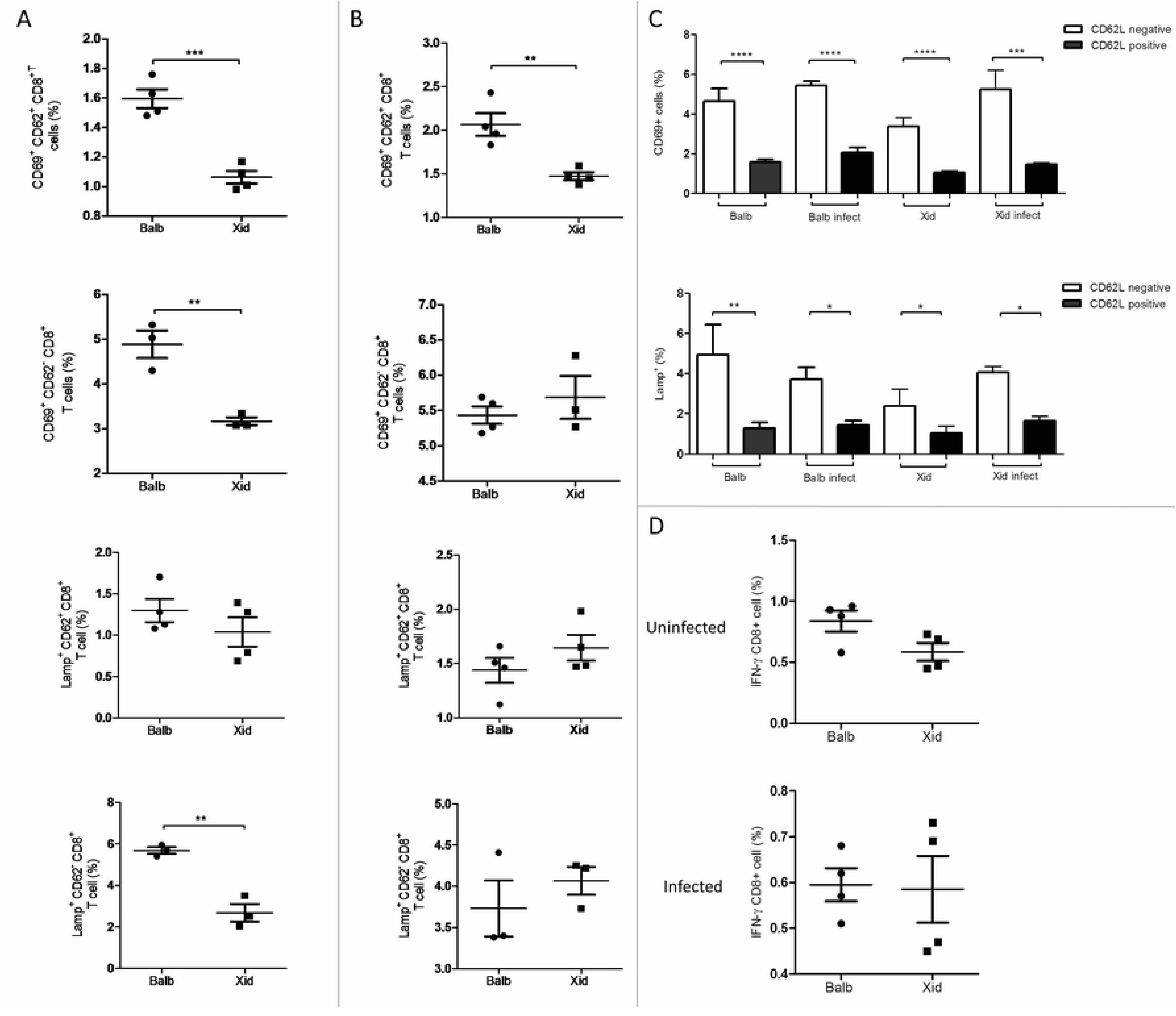
Expression of surface activation molecules and intracellular interferon in CD8 T lymphocytes from Balb (WT) and Xid mice not infected (Uninfected) or infected (Infected) with *E. cuniculi* spores. (A) Analysis of the expression of the selectins CD62L, Lamp 1, and CD69 on the surface of CD8^+^ T lymphocytes from the spleen of uninfected Balb/c and Xid mice that were co-cultured with *E. cuniculi* infected macrophages. (B) Analysis of the expression of the selectins CD62L, Lamp 1, and CD69 on the surface of CD8+ T lymphocytes from the spleen of Balb/c mice previously infected with *E. cuniculi* and co-cultured with *E. cuniculi*-infected macrophages. (C) The expression of selectins CD62L (positive or negative) x CD69 or CD62L (positive or negative) x Lamp 1 on the surface of CD8^+^ T lymphocytes from the spleen of uninfected Balb/c or Xid mice and infected Balb or Xid mice. (D) The intracellular expression of INF-γ in CD8*^+^* T lymphocytes from the spleen of uninfected Balb/c and Xid mice or Balb/c and Xid mice previously infected with *E. cuniculi*, all of which were co-cultured with macrophages infected with *E. cuniculi*. Statistical analysis: paired t-tests were conducted; *p < 0.05, **p < 0.01, ***p < 0.001, and ****p < 0.0001

## Discussion

Several researchers have found that B-1 cells participate in the response mediated by T lymphocytes, for example, in immediate or delayed hypersensitivity reaction [27, 28] and allograft rejection [25]. B-1 cells can also promote the activation of T lymphocytes and the production of IFN-γ by adoptively transferred T lymphocytes [26]. Our group found that intraperitoneal infection of Xid mice with *E. cuniculi* resulted in an increase in CD8^+^ T lymphocytes in the spleen and peritoneum, although these mice showed a greater range of clinical signs and fungal burden, which indicated lower resistance to infection [21, 23]. In Xid mice infected orally with *E. cuniculi*, the decrease in the number of CD8^+^ T cells in the spleen and peritoneum was associated with a decrease in the content of serum IFN-γ. These findings indicated that the resistance to infection was low [22].

In this study, we found that CD8 T lymphocytes from WT (naïve) mice showed high proliferative capacity, a high percentage of CD69 activation molecules, a low mortality rate, and high microbicidal activity when cultured with macrophages. These results suggested that B-1 cells present in Balb mice potentiated the cytotoxic activity of lymphocytes. However, CD8 T cells from Xid mice (deficient in B-1 cells) proliferated less. They exhibited lower expression of activation molecules and lower microbicidal activity of macrophages in co-cultures. Our findings suggested that the presence of B-1 cells (in Balb mice) or their deficiency (in Xid mice) altered the cytotoxic function of CD8 T lymphocytes. These changes could explained why Xid mice showed low resistance to *E. cuniculi* infection, although the population of CD8 T cells was quite large. Our results were similar to those of studies in which B1-cells were found to potentiate the activity of CD8 T lymphocytes [25, 26, 27, 28]. Thus, the relationship between lymphocytes/B-1 cells/macrophages might be linked to functional factors involving complex signaling pathways.

The expression of CD69 in infiltrating lymphocytes in inflamed tissues is a marker of different signaling pathways, which can regulate tissue retention, metabolism, and their activated phenotype [29]. CD69 expression is rapidly induced on the surface of T lymphocytes after the interaction between TCR and CD3, and was responsible for activates cytokines, polyclonal antibodies, and mitogens [30, 31]. Consistent with this information, we found that CD8 T lymphocytes from WT mice showed higher expression of CD69 and were associated with an increase in the microbicidal activity of macrophages infected with *E. cuniculi*. The proliferation of CD8 T lymphocytes in this group of mice was also higher, probably due a mitogenic stimulus. However, the reduction in the microbicidal activity of macrophages co-cultured with CD8^+^CD69^low^ T lymphocytes was determined by the high proliferation of spores recovered from these assays. These findings confirmed that a positive relationship occurred between CD69 expression in CD8 T cells and macrophage stimulation, as recorded in tumor macrophages (TAM) [32]. The CD8 T lymphocytes obtained from infected Xid mice (primed) with *E. cuniculi* showed a higher mortality rate due to necrosis and less for apoptosis. A similar finding was recorded for macrophages co-cultured with these lymphocytes. In another study, we showed that macrophages infected with *E. cuniculi* have a higher rate of apoptosis, which matched the findings described here [33]. The natural response of an infected cell is to undergo apoptosis to prevent the pathogen from multiplying and spreading. However, apoptosis can act as a mechanism to evade the immune response; for example, *Encephalitozoon* can suppress apoptosis, as found in Vero cells, by inhibiting the cleavage of caspase-3, phosphorylation, and translocation of p53 [34]. We also found that *E. cuniculi* can exploit efferocytosis to enter the host cell, multiply, and spread, as well as, modulate to a less inflamed environment. These changes act as a favorable mechanism for evading immunity [33].

We also found that lymphocytes from previously infected Xid mice showed higher expression of CD107, demonstrating their ability to function against *E. cuniculi.* Lamp-1 (lysosome-associated membrane protein CD107a) is predominantly expressed intracellularly, but after activation, it can be expressed on the cell surface. LAMP-1 comprises 50% of all lysosomal membrane proteins and is widely used as a cell surface marker of lymphocyte activation and degranulation. Additionally, LAMP-1 is a physiologically essential protein involved in stabilizing lysosomes and regulating autophagy to prevent embryonic lethality [35, 36, 37]. The expression of CD107 is associated with a decrease in the expression of CD62L. Also, the elimination of CD62L and changes in the cytoskeleton are associated with an increase in the lytic activity of T cells [38]. We hypothesized that higher expression of CD107 and proliferative activity might explain the high microbicidal activity exerted by macrophages co-cultured with Xid lymphocytes previously infected with *E. cuniculi*, considering that memory CD8 T cells have greater potential for division, high chance of survival, and low effector activity. In conclusion, our results suggest that the absence of B-1 cells in Xid mice might be associated with the lower expression of activating molecules in CD8^+^ T lymphocytes and their cytotoxic activity. However, after a previous infection with *E. cuniculi*, CD8 T lymphocytes were more effective in killing macrophages infected with *E. cuniculi*.

## Data availability

Data will be made available on reasonable request.

